# Anterior pituitary gland volume mediates associations between pubertal hormones and changes in transdiagnostic symptoms in youth

**DOI:** 10.1101/2024.05.17.594766

**Authors:** Giorgia Picci, Nathan M. Petro, Chloe C. Casagrande, Lauren R. Ott, Hannah J. Okelberry, Danielle L. Rice, Anna T. Coutant, Grace C. Ende, Erica L. Steiner, Yu-Ping Wang, Julia M. Stephen, Vince D. Calhoun, Tony W. Wilson

**Author notes:** **Corresponding author:** Giorgia Picci, PhD Research Scientist, Laboratory Director Institute for Human Neuroscience Boys Town National Research Hospital 14090 Mother Teresa Lane Boys Town, NE, USA.

## Abstract

The pituitary gland (PG) plays a central role in the production and secretion of pubertal hormones, with documented links to the emergence and increase in mental health symptoms known to occur during adolescence. Although much of the literature has focused on examining whole PG volume, recent findings suggest that there are associations among pubertal hormone levels, including dehydroepiandrosterone (DHEA), subregions of the PG, and elevated mental health symptoms (e.g., internalizing symptoms) during adolescence. Surprisingly, studies have not yet examined associations among these factors and increasing transdiagnostic symptomology, despite DHEA being a primary output of the anterior PG. Therefore, the current study sought to fill this gap by examining whether anterior PG volume specifically mediates associations between DHEA levels and changes in dysregulation symptoms in an adolescent sample (*N* = 114, 9 – 17 years, M_age_ = 12.87, SD = 1.88). Following manual tracing of the anterior and posterior PG, structural equation modeling revealed that greater anterior, not posterior, PG volume mediated the association between greater DHEA levels and increasing dysregulation symptoms across time, controlling for baseline dysregulation symptom levels. These results suggest specificity in the role of the anterior PG in adrenarcheal processes that may confer risk for psychopathology during adolescence. This work not only highlights the importance of separately tracing the anterior and posterior PG, but also suggests that transdiagnostic factors like dysregulation are useful in parsing hormone-related increases in mental health symptoms in youth.

## Introduction

The pituitary gland (PG) has been highlighted as the nexus of neuroendocrine regulation and stress responsivity, given its primary role in hypothalamic-pituitary-adrenal (HPA) axis function. Clear links have been made between morphometric estimates of the PG and responses to stressful experiences, as well as mental health functioning during development. Although the majority of studies have focused on whole pituitary volumes (Ganella et al., 2015; Kaess et al., 2013; Murray et al., 2016; Whittle et al., 2012; Zipursky et al., 2011), emerging work has suggested that the anterior PG, in particular, is highly relevant to puberty-related responses to early life stress and mental health symptoms that occur within the adolescent window (Díaz-Arteche et al., 2021; Farrow et al., 2020; Picci et al., 2023a). That is, the PG consists of two separable lobes that have dissociable structural and functional features. The anterior PG is primarily involved in the production and release of a variety of hormones (Kaiser and Ho, 2016), including dehydroepiandrosterone (DHEA), which is a key hormone in pubertal development and the focus of the current investigation. Indeed, with adrenarche, the anterior PG plays an increasing role in the influx of DHEA, with concomitant volumetric increases in whole PG volume during this early pubertal event (Murray et al., 2016). Conversely, the posterior PG is largely responsible for the storage and release of antidiuretic hormone (i.e., vasopressin) and oxytocin, which are produced by the hypothalamus (Leng et al., 2015). Thus, the anterior PG, in particular, is of interest because its morphology may offer insights into the puberty-related emergence of mental health symptoms. Importantly, far less is known about how adrenarche and its associated hormones are related to adolescent mental health (Byrne et al., 2017); though such studies could offer foundational knowledge about how even the earliest pubertal events (i.e., before the major physical changes associated with puberty) set the stage for adolescent mental health functioning. In other words, because adrenarche most often occurs before gonadarche, earlier detection of neurally-mediated adrenarche-related alterations in symptomology could yield greater prediction of who is most vulnerable to developing mental health disorders in the long term.

Specific links between adrenarche (e.g., DHEA levels), anterior PG volume, and mental health symptoms in youth have yet to be properly documented, especially in longitudinal samples. Generally, existing work has largely focused on whole PG volumes. Although mixed (Díaz-Arteche et al., 2021; Farrow et al., 2020; Ganella et al., 2015), there are reports that larger whole PG volume is related to subclinical and clinical levels of mental health symptoms. For example, longitudinal work has shown that larger PG volume prospectively predicts increases in internalizing symptoms during the pubertal window (Zipursky et al., 2011). Similarly, larger PG volume was found to mediate associations between elevated depressive symptomology and earlier puberty onset (Whittle et al., 2012). This corroborates earlier studies showing that youth diagnosed with unipolar and bipolar depression had larger PG volume than controls (MacMaster et al., 2008). More recently, work has highlighted the contribution of early life stress in accelerating PG growth (Ganella et al., 2015; Kaess et al., 2018) or promoting larger PG volumes into adulthood (Klinger-König et al., 2023). That is, in a population-based study of adults (N > 2800), PG volumes were related to psychopathology symptoms, which were further amplified by early life stress (Klinger-König et al., 2023). Discrepancies in existing findings include that some of the accelerated growth findings are only evident in girls (Ganella et al., 2015; Kaess et al., 2018) and that in some cases, PG volume does not relate to symptomology levels (Ganella et al., 2015) nor early life stress (Schär et al., 2022).

Existing work has also characterized how pubertal hormone levels are related to PG volume during the pubertal transition, and how this may correspond with the emergence of psychopathology. In one study, salivary testosterone, DHEA, and DHEA-S predicted increases in PG volume across time, irrespective of age (Whittle et al., 2020). Notably, this pattern of growth attributable to hormones was stronger for girls than boys. PG volume has also been shown to mediate associations between higher DHEA/DHEA-S levels and anxiety symptoms in youth (Murray et al., 2016). In contrast, other work has shown that youth with elevated trauma-related anxiety symptoms exhibit lower DHEA levels, which in turn predict smaller anterior PG volumes specifically (Picci et al., 2023a). Although contrary to some developmental work, these findings corroborate other literature showing both neuroprotective effects of elevated DHEA and systematically lower levels of DHEA in those with internalizing disorders (Farooqi et al., 2018; Mocking et al., 2015). This prior work motivates a more targeted, longitudinal investigation of whether anterior PG volume specifically mediates associations between DHEA levels and symptomology across adolescence.

Although foundational, extant literature has overwhelmingly examined whole PG volume, with more limited study of whether and to what extent the anterior PG is a key arbiter in emergent symptomology in adolescence. Some studies have begun to examine the anterior and posterior pituitary lobes separately, which may help to disentangle some of the discrepant findings by enhancing overall sensitivity to detect effects. Given its specific role in stress responsivity and pubertal hormone output (Le Tissier et al., 2012; Martí and Armario, 1998), it may be the case that there are specific stress- and hormone-related patterns of variability evident in the anterior versus the posterior lobe of the PG. The sparse findings that are available suggest that there is utility in separating the anterior and posterior lobes, as lobe specific effects have been uncovered (Díaz-Arteche et al., 2021; Farrow et al., 2020; Picci et al., 2023a). That is, two prior studies emphasize the role that early life stress may have in propagating risk for enlarged anterior PG volumes specifically, although neither of these studies found prospective links with psychopathology, and they did not include pubertal hormones (Díaz-Arteche et al., 2021; Farrow et al., 2020). Taken together, the current literature suggests that the anterior pituitary, in particular, may be a viable target for understanding associations among pubertal hormones, a stress sensitive neural region, and emergent psychopathology. Therefore, the current study is among the first longitudinal investigations to examine how the morphometry of the anterior PG may mediate associations between adrenarcheal hormone levels and changes in symptomology during adolescence.

Given that previous research has not yet shown that DHEA levels and pituitary volume amplify risk for specific diagnostic categories, the current study employed a more transdiagnostic approach. We built off of previous work highlighting that dysregulation is a potent predictor of long-term psychopathology diagnoses and comorbidities (Althoff and Ametti, 2021), as well as changes in key neural networks (Picci et al., 2023b). Dysregulation is often defined as difficulties in regulating affect, behavior, and cognition (ABCs), with symptoms peaking during early adolescence (Deutz et al., 2018). As a symptom profile, it does not completely fit into either internalizing or externalizing categories, but instead, has features of both. In the Child Behavior Checklist Dysregulation Profile (CBCL-DP) employed here, dysregulation consists of the anxiety/depression, attention problems, and aggression subscales. Thus, the current study examined the extent to which DHEA levels (adrenarche) were indirectly associated with changes in dysregulation symptomology during adolescence, via anterior PG volume. Building upon previous literature, we predicted that enlarged anterior (but not posterior) PG would mediate the relationship between higher DHEA levels and increases in dysregulation symptoms across time.

## Methods

A final sample of 114 healthy children and adolescents ages 9–17 years-old completed a structural MRI scan and a hormonal assay as part of the Developmental Chronnecto-Genomics study (mean_age_= 12.87, SD = 1.88; 58 females; Stephen et al., 2021). The study was multisite, with 44 participants recruited at the Boys Town National Research Hospital site in Omaha and 70 participants from the Mind Research Network (MRN) in Albuquerque, NM, USA. Participants were invited back to participate annually for three years. Participants’ symptomology levels were collected via caregiver report annually (see *Child Behavioral Checklist* section). In the present study, we examined structural MRI data and hormonal assays from the same time point and change in symptomology across two timepoints. Specifically, we examined change in symptomology from the scan/hormone assay session to the participant’s next visit approximately 1 year later. The initial sample included 153 participants with a T1-weighted image and a usable hormonal assay; 16 participants (14%) were excluded due to excessive motion and/or artifacts in their structural MRI data and 23 were excluded for missing two consecutive timepoints of CBCL data. Inclusion criteria included English as a primary language, age, and participant and parent willingness to assent/consent, respectively. Exclusion criteria determined via parent report were as follows: history of developmental delays and/or diagnosed psychiatric disorders, history of neurological disorders (e.g., epilepsy), history of concussion or head injury, pregnancy, prenatal exposure to drugs, use of medications known to affect brain function, and MRI contraindications (e.g., orthodontia, metallic foreign bodies). All parents and youth provided written consent or assent, respectively, prior to participating in the study. The appropriate institutional review boards for both study sites approved all study procedures.

### Structural Neuroimaging Acquisition & Processing

Participants underwent a structural T1-weighted MRI with either a Siemens 3T Skyra scanner at the Boys Town National Research Hospital study site (N=44) or a Siemens 3T TIM Trio at the MRN site (N=70). MR images at both data collection sites were acquired with a 32-channel head coil and an isotropic MPRAGE sequence with the following parameters: TR=2400 milliseconds; TE=1.94 milliseconds; flip angle=8°; field of view=256 mm; slice thickness=1 mm; base resolution=256; 192 slices; voxel size=1×1×1 mm. The images of all participants were processed using FreeSurfer version 6 (http://surfer.nmr.mgh.harvard.edu) to extract total intracranial volume estimates. We followed the ENIGMA protocol for quality assurance, including performing visual checks on all images (http://enigma.usc.edu/protocols/imaging-protocols) and checking for motion artifacts.

### Pituitary Tracing

Tracing methods for this study have been reported previously (see Picci et al., 2023a). Briefly, the anterior and posterior lobes of the pituitary gland were manually traced using the 3D Slicer software (v4.11, www.slicer.org; Fedorov et al., 2012). Prior work has demonstrated that the pituitary is optimally visualized in the coronal and sagittal planes of T1-weighted MR images (Díaz-Arteche et al., 2021; Farrow et al., 2020; Lorenzetti et al., 2009; Whittle et al., 2012). The pituitary was defined based upon previously published tracing techniques (Farrow et al., 2020). Several boundaries and structures were visually inspected by the tracers to determine the boundaries of the PG (Figure 1, adapted from Picci et al., 2023). That is, the diaphragm sellae and lateral ventricles were located superiorly, the sphenoid sinus was located inferiorly and bilaterally, and the internal carotid arteries were located bilaterally, all in the coronal view. Contrast differences in the sagittal view were used to determine the boundary between the anterior and posterior lobes of the PG, which appear darker and lighter, respectively (Figure 1). Estimates for the anterior and posterior lobe volumes were computed by summing all the voxels selected by tracers across all slices. The full sample was split in half and traced by 2 pairs of trained tracers (i.e., LO-NP, GP-CC were paired tracers). Each tracer traced both the anterior and posterior lobes (for example tracing, see Figure 2). Intraclass coefficients (ICCs) with absolute agreement were calculated for anterior and posterior traces, and were .96-.97 (LO-NP) and .93-.95 (GP-CC), respectively.

**Figure 1.**
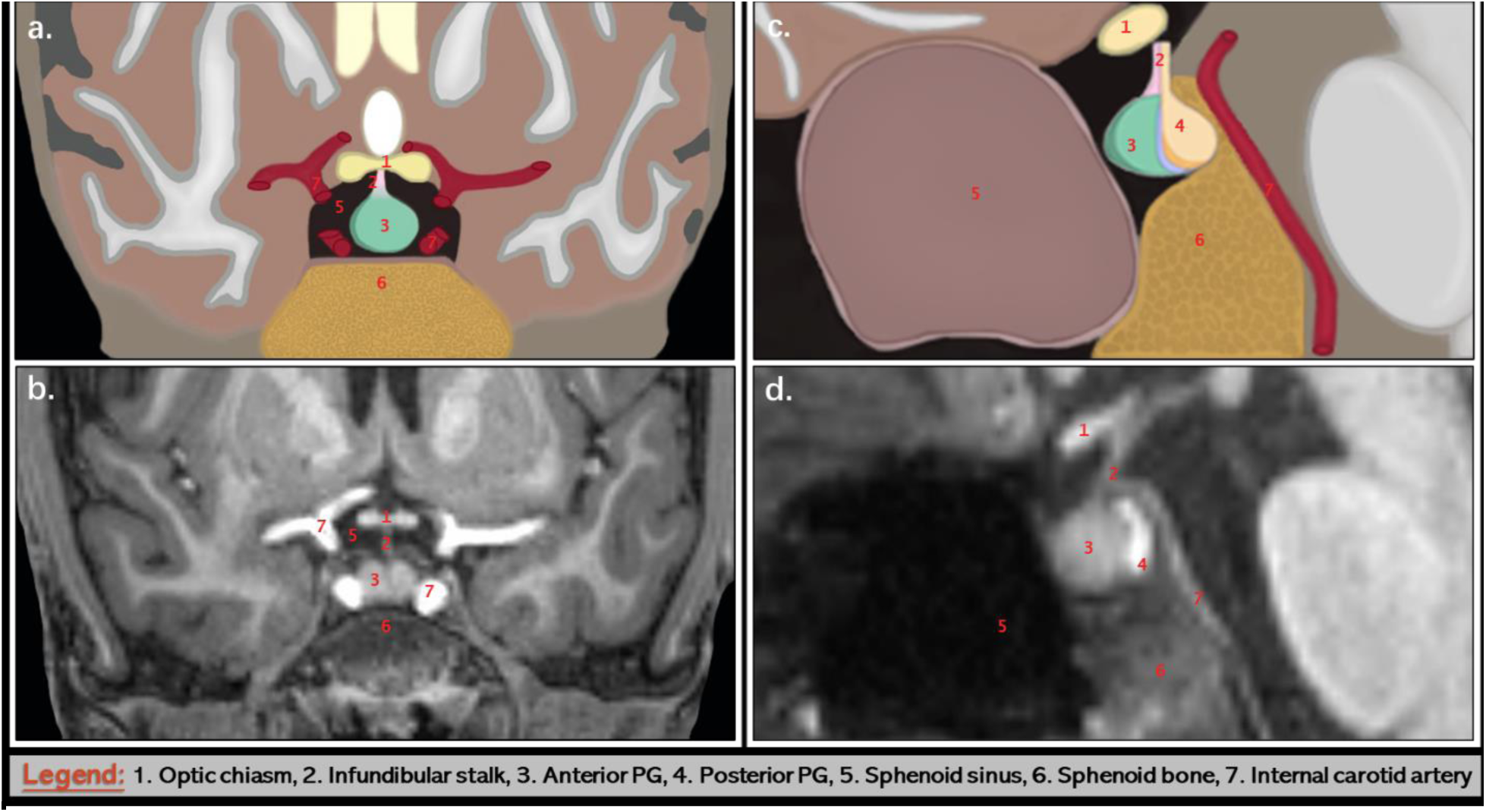
Anatomical boundaries used to trace both lobes of the pituitary gland. (a) and (c) are coronal and sagittal schematic depictions of the T1-weighted slice images from a representative participant in (b) and (d). The red numbers denote the anterior and posterior lobes of the pituitary gland (PG), as well as the structures and anatomical boundaries surrounding them. Each of the numbered structures were used to manually trace the anterior and posterior PG. The legend (bottom of figure) provides the anatomical structures that correspond to each of the red numbers in a-d.

**Figure 2.**
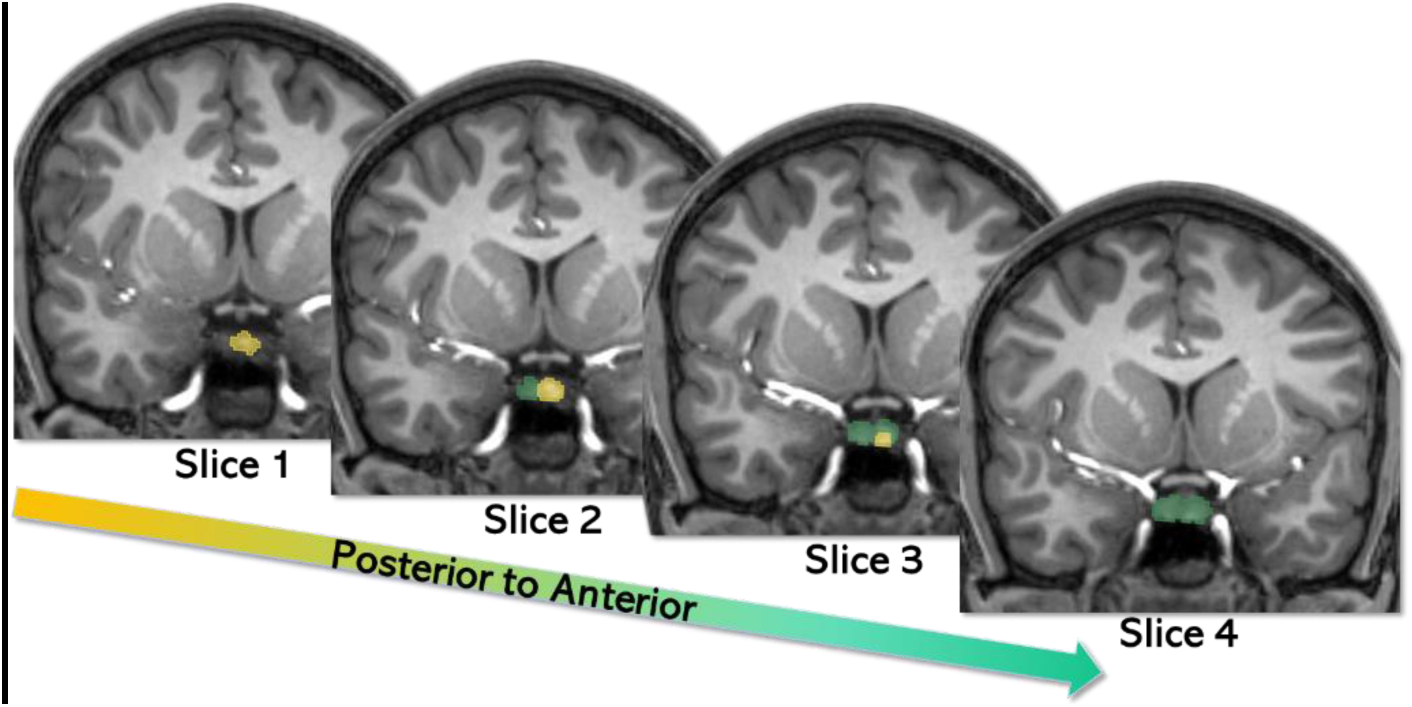
Slice-by-slice example of a manually traced pituitary gland. A set of T1-weighted images in the coronal view from a representative participant is shown across multiple slices proceeding from posterior to anterior. The anterior pituitary gland (PG) is shown in teal/green and the posterior PG shown in yellow.

### Child Behavior Checklist

During two of the study visits, a caregiver completed the Child Behavior Checklist (CBCL, (Achenbach et al., 2001)) to assess their child’s dysregulation behaviors over the past 6 months. The dysregulation profile is a summed score of the attention, aggression, and anxious/depressed subscales (Holtmann et al., 2011). Scores from the participant’s initial visit where they had both a usable T1-weighted structural scan and hormonal assays served as the baseline measure of dysregulation symptoms. Scores from the participant’s next visit, approximately 1 year later, served as time point 2. Raw scores were used in our models, as both sex and age were controlled for in our models. A difference score was calculated for each individual to assess changes in dysregulation across time (i.e., Time 2 symptoms – Time 1 symptoms). Therefore, positive change scores indicate increasing dysregulation across time, whereas negative change scores indicate decreasing dysregulation symptoms.

### Hormonal Assays

At least 2.0 ml of whole unstimulated saliva was collected from each participant. Specifically, children were asked to passively drool into an Oragene DISCOVER (OGR-500; www.dnagenotek.com) collection tube until liquid saliva (not bubbles) exceeded the fill line indicated on the tube. Participants were instructed to refrain from consuming any food, liquids, or chewing gum for at least an hour before providing the saliva sample, and generally completed the study in the afternoon. Prior to the release of the protease inhibitors for long-term storage, a single-channel pipette was used to extract 0.5 ml from the collection tube, which was immediately transferred into a labeled micro-centrifuge tube and placed in a -20C freezer for storage. All samples were assayed together with duplicate testing using a commercially-available assay kit for salivary DHEA. The assay kit had a sensitivity of 5 pg/mL, with a range of 10.2-1000 pg/mL. The intra- and inter-assay coefficients of variation were 7.95% and 10.17%, respectively. The averages of duplicate tests were used for analyses in the present study. A natural log transform was applied to DHEA measurements to account for skewness of the raw data (before transform: skewness=3.32, kurtosis=13.34; after transform: skewness=0.36, kurtosis=-0.04).

### Data Analytic Plan

We first ran descriptive statistics on all study variables of interest and demographics. Variables entered into subsequent models were examined for violations of normality (i.e., skewness and kurtosis) and were transformed according to their distribution type (e.g., positive versus negative skew). Next, we iteratively fit structural equation models to estimate 1) whether anterior and/or posterior PG volumes mediated associations between DHEA levels and changes in dysregulation symptoms across time (see Figure 3 for model). Note that we also ran models separately by sex to determine sex-specific patterns, given known sex differences in pubertal timing and mental health symptomology; these results are reported in the Supplement.

**Figure 3.**
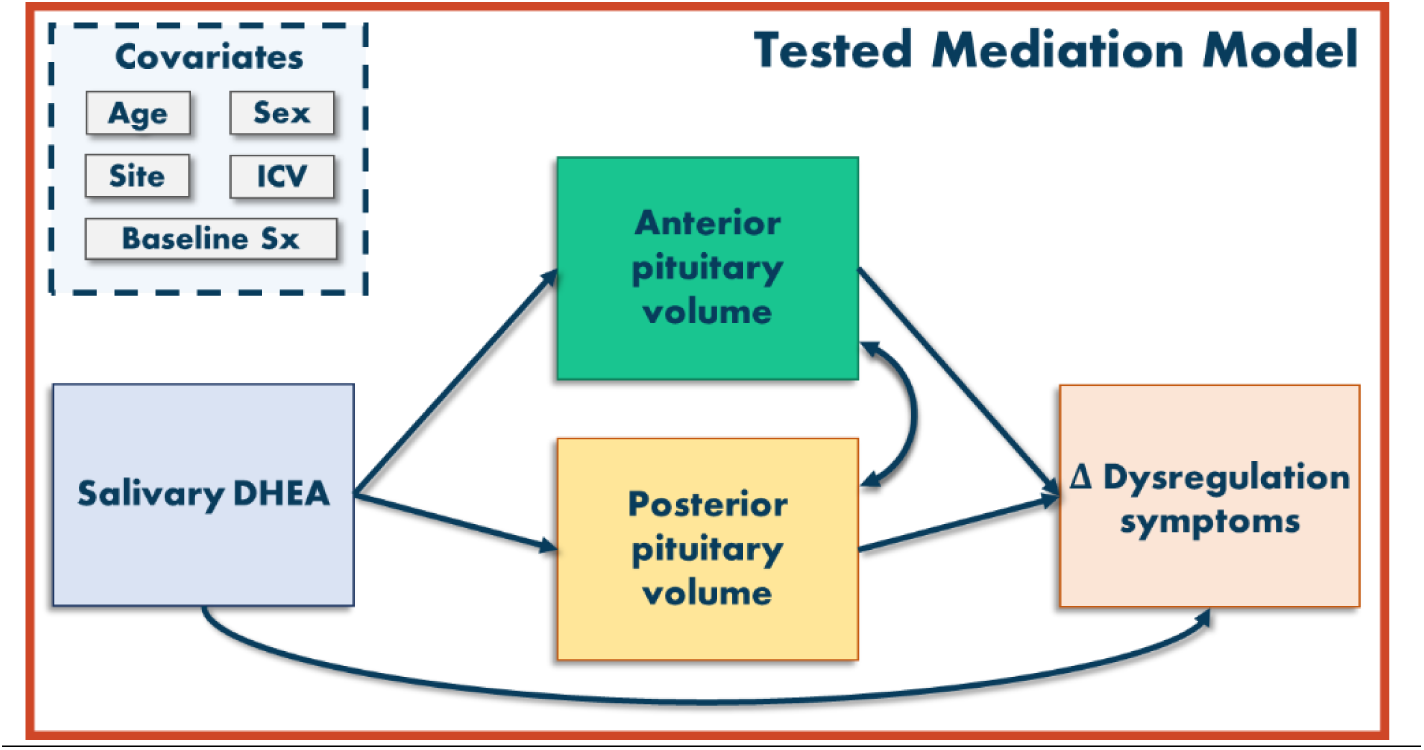
Full structural equation model used in the current study. The mediation model tested whether anterior and/or posterior pituitary gland (PG) volume mediated the association between salivary DHEA levels and changes in dysregulation symptoms across two time points measured via the CBCL. Dysregulation symptom change scores were computed by subtracted time 1 scores from time 2 scores; larger values indicate increases in symptoms across time. All study variables were regressed on sex, age, study site, and baseline dysregulation symptoms. PG variables were regressed on ICV. ICV = intracranial volume, baseline Sx = baseline dysregulation symptoms, Δ Dysregulation symptoms = dysregulation change scores.

Base models were fit to examine the effect of independent variables on the dependent variables. Next, covariates of no interest were introduced into the model, which included sex, age, study site, baseline levels of dysregulation symptoms, and total intracranial volume (ICV). Note that ICV was regressed only on anterior and posterior PG volumes. After introducing the covariates, models were fit to examine the mediating effect of anterior and posterior PG volume on the association between DHEA levels and symptomology levels (i.e., indirect effects of PG volume via DHEA on symptomology). All mediation models were bootstrapped with 1000 iterations with bias-corrected bootstrapping to test for significance of the indirect associations based upon the 95% confidence intervals (MacKinnon et al., 2004). Unstandardized estimates of individual paths were examined for the directionality of relations with an *a priori p*-value significance threshold of *p* < .05. Finally, PG volume estimates (i.e., anterior and posterior lobe volumes) were permitted to freely correlate and all parameters were freely estimated. All models were tested in Mplus (v7.4).

We examined the goodness of fit for both models using standard criteria (Hu and Bentler, 1999). Specifically, we evaluated models for the root mean square error of approximation (RMSEA) < 0.06, standardized root mean square residual (SRMR) < 0.08, and comparative fit index (CFI) > 0.95. We also examined the χ^2^ test of model fit, where a nonsignificant result indicates good model fit.

## Results

### Sample Demographics

Descriptive statistics and demographic variables for the final sample (N=114) are reported in Table 1. Correlations among study variables of interest are reported in Table S1. There were study site differences in ethnicity such that the MRN site included more Latinx/Hispanic participants compared to the Boys Town National Research Hospital (BTNRH) site (all *p* < .05, Table 1). In addition, there were site differences in anterior and posterior PG volumes, whereby the BTNRH site had significantly larger volumes for both segmentations (all *p* < .05, Table 1). There were no other site differences in participant distributions of age, sex, race, hormone levels, and symptomology. Crucially, study site was included in all models as a covariate to account for these initial differences.

**Table 1.** Demographics and study site comparisons.

| Variable | Full Sample<br>(n = 114) |  | Site 1<br>(BTNRH; n = 44) |  | Site 2<br>(MRN; n = 70) |  | Comparison |  |
| --- | --- | --- | --- | --- | --- | --- | --- | --- |
| | n | % | n | % | n | % | $\chi^2$ | p |
| **Sex, female** | 58 | 51% | 26 | 59% | 32 | 46% | 1.93 | .16 |
| **Race** |  |  |  |  |  |  | 6.03 | .20 |
| AI/AN | 5 | 4% | - | - | 5 | 7% |  |  |
| Asian | 1 | 1% | 1 | 2% | - | - |  |  |
| B/AA | 5 | 4% | 3 | 7% | 2 | 3% |  |  |
| White | 96 | 86% | 37 | 84% | 59 | 84% |  |  |
| More than one race | 4 | 3% | 1 | 2% | 3 | 4% |  |  |
| Not reported | 3 | 2% | 2 | 5% | 1 | 2% |  |  |
| **Ethnicity** |  |  |  |  |  |  | 9.56 | **<.05** |
| Latinx/Hispanic | 25 | 22% | 3 | 7% | 22 | 31% |  |  |
| Not Latinx/Hispanic | 89 | 78% | 41 | 93% | 48 | 69% |  |  |
| Not reported |  |  |  |  |  |  |  |  |
|  | Mean (SD) | Range | Mean (SD) | Range | Mean (SD) | Range | t | p |
| **Age (years)** | 12.87 (1.88) | 9 – 17 | 12.82 (1.61) | 9 – 15 | 12.89 (2.05) | 9 – 17 | -0.20 | .84 |
| **DHEA** | 53.09 (63.90) | 5.15 – 417.99 | 44.26 (49.86) | 5.15 – 302.48 | 58.63 (71.11) | 6.58 – 417.99 | -1.17 | .24 |
| **Dysregulation symptoms** | 6.06 (6.07) | 0 – 30 | 5.45 (5.41) | 0 – 20 | 6.45 (6.45) | 0 – 30 | -0.83 | .41 |
| **$\Delta$ Dysregulation** | 0.75 (5.02) | -12 – 21 | 0.52 (4.24) | -8 – 13 | 0.90 (5.49) | -12 – 21 | -0.33 | .74 |
| **Anterior pituitary volume** | 404.04 (155.87) | 87 – 851 | 442.66 (158.96) | 158 – 844 | 379.76 (149.98) | 87 – 851 | 2.13 | **<.05** |
| **Posterior pituitary volume** | 87.57 (35.84) | 29 – 215 | 112.28 (33.88) | 65 – 215 | 72.04 (27.47) | 29 – 166 | 6.95 | **<.001** |

**Note.** AI/AN = American Indian/Alaska Native, B/AA = Black, African American. Brain measures are reported in volume mm<sup>3</sup>. Age is reported in years, at the time of the MRI scan. DHEA is reported in picograms per milliliter (pg/mL). Dysregulation symptoms are all raw scores from the Child Behavior Checklist. Dysregulation change scores ( $\Delta$ ) were calculated as follows: T2 – T1 dysregulation symptoms; positive scores indicate increasing symptom severity across time. To evaluate potential study site effects, $\chi^2$ analyses were conducted for binary variables. Significantly different p-values are bolded.

### Mediation Model Results

The mediation model had excellent fit [*χ*^2^(2) = 1.34, *p* = .51; RMSEA < 0.01; SRMR = 0.01; CFI = 1.00]. Table 2 reports all direct and indirect effects and all path estimates are reported in Table S2. In terms of the mediation results, anterior PG volume mediated the association between DHEA and changes in dysregulation symptoms across time (β = 0.10, *b* = 0.08; 95% CI[0.02, 0.19]; Figure 4). There was a direct effect between DHEA levels and changes in dysregulation symptoms (β = -0.31, *b* = -0.25, *p* = .02). Participants with higher DHEA levels had larger anterior PG volumes (β = 0.28, *b =* 0.51*, p* < .001), which in turn predicted greater increases in dysregulation symptoms across time (β = 0.35, *b =* 0.16*, p* = .03).

**Figure 4.**
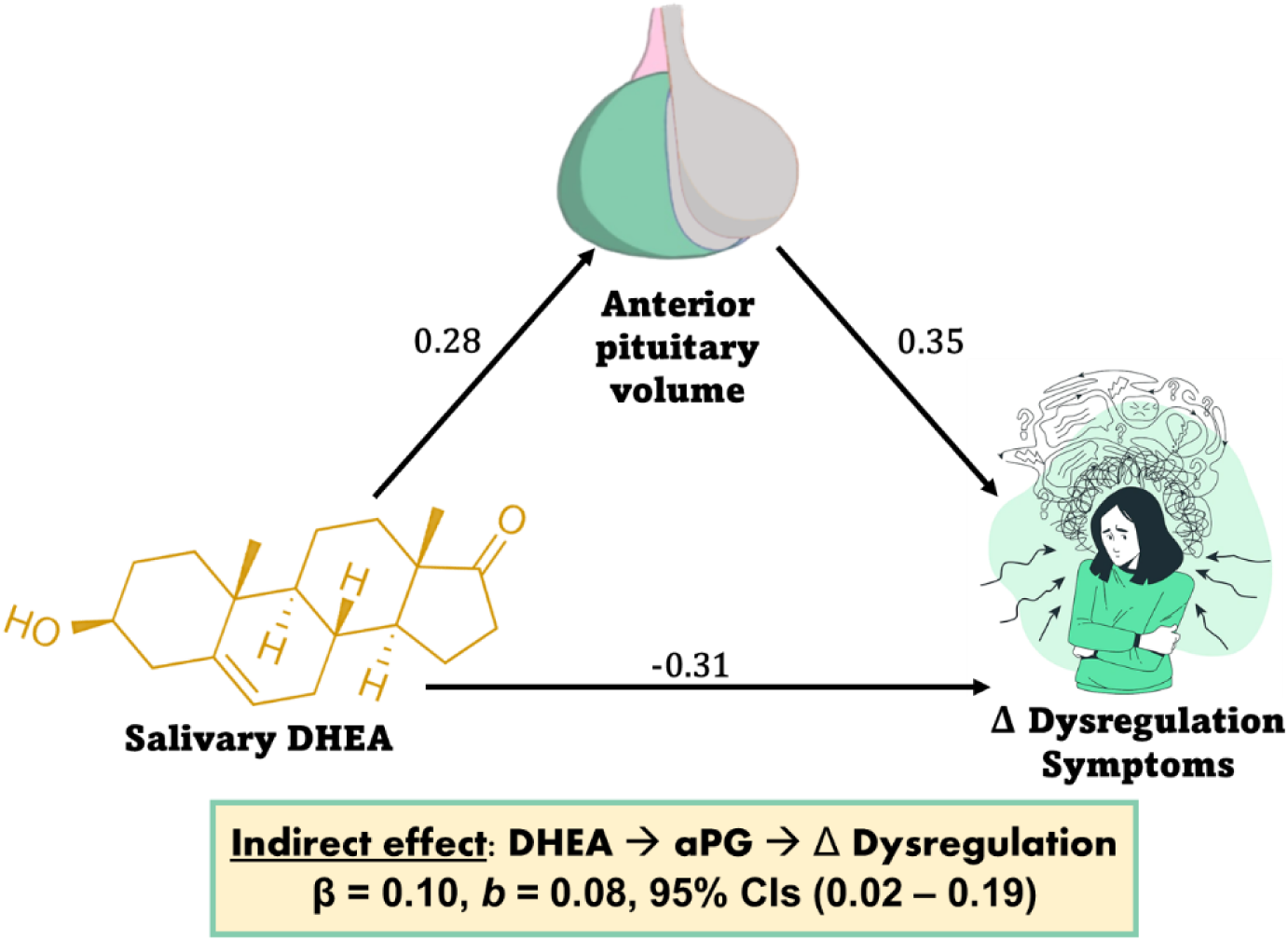
Significant indirect effect of anterior pituitary volume on associations between DHEA and changes in dysregulation. Higher DHEA levels were related to larger anterior pituitary volume, which in turn predicted greater increases in dysregulation symptoms across time. There was also a direct relationship between DHEA and symptoms such that greater DHEA levels related to fewer symptoms. Standardized estimates are shown. Model fit was excellent.

**Table 2.** Direct and indirect effect estimates for mediation model.

| Indirect Effects |  | Estimates |  | 95% CIs |  |
| --- | --- | --- | --- | --- | --- |
| Mediator | Outcome | $\beta$ | $b$ | Lower | Upper |
| **Anterior pituitary** | **$\Delta$ Dysregulation** | 0.10 | 0.08 | **0.02** | **0.19** |
| Posterior pituitary | $\Delta$ Dysregulation | 0.005 | 0.004 | -0.01 | 0.07 |

| Direct Effects |  | Estimates |  | 95% CIs |  |
| --- | --- | --- | --- | --- | --- |
| | | $\beta$ | $b$ | Lower | Upper |
| **DHEA → $\Delta$ Dysregulation** |  | -0.31 | -0.25 | **-0.47** | **-0.03** |

Note. Significant results are bolded based upon bootstrapped 95% confidence intervals for indirect effects and $p < .05$ for direct effects.

Mediation effects were specific to the anterior PG such that the posterior PG did not mediate associations between DHEA levels and changes in dysregulation symptoms (β = 0.005, *b* = 0.004; 95% CI[-0.01, 0.07]). There were no associations between DHEA levels and posterior PG volume (β = -0.10, *b* = -0.04, *p* = .30), nor were there associations between posterior PG and changes in dysregulation (β = -0.05, *b* = -0.10, *p* = .72).

Other notable effects in the model include sex-specific effects on the anterior (β = 0.38, *b* = 1.23, *p* < .001), but not the posterior PG (β = -0.05, *b* = -0.03, *p* = .62). Specifically, girls had significantly larger anterior PG compared to boys. In addition, there were no sex differences in DHEA levels (β = 0.12, *b*=-0.21, *p* = .14) nor changes in dysregulation (β = -0.03, *b* = -0.04, *p* = .81). Finally, age was associated with increasing DHEA (β = 0.53, *b* = 0.24, *p* < .001), anterior pituitary volume (β = 0.40, *b* = 0.34, *p* < .001), and posterior pituitary volume (β = 0.28, *b* = 0.05, *p* = .002), but not changes in dysregulation symptoms (β = 0.08, *b* = 0.03, *p* = .61).

## Discussion

The current study is among the first to establish longitudinal associations between pubertal hormones, anterior PG volume, and changes in transdiagnostic symptomology during the pivotal adolescent period. Here, we report that greater anterior PG volume mediates the relationship between higher DHEA levels at baseline and increases in dysregulation symptomology across time. Crucially, these findings were only found in the anterior lobe of the PG, not the posterior lobe, suggesting specificity of effects in a region integral to HPA-axis functioning, which is consistent with extant literature (Farrow et al., 2020; Picci et al., 2023a). There were also no direct relationships between posterior PG volume and DHEA levels nor changes in dysregulation symptomology, further suggesting that the anterior PG is uniquely sensitive to pubertal hormones and is consequentially linked to mental health during the pubertal window. Although contrary to expectations, the direct relationship between greater DHEA and decreasing dysregulation symptomology adds to the literature showing neuroprotective effects of DHEA during development (Farooqi et al., 2018) and is consistent with systematically lower levels of DHEA in adults with internalizing disorders (Mocking et al., 2015).

The mediation finding reported here is largely consistent with prior cross-sectional and longitudinal literature documenting associations among different metrics of puberty, PG volume, and symptoms during adolescence (Murray et al., 2016; Whittle et al., 2012; Zipursky et al., 2011). Specifically, previous work has shown a comparable mediation effect with whole PG volume, pubertal timing, and depressive symptoms across time (Whittle et al., 2012). The present study builds upon prior findings by documenting greater specificity of these effects in the anterior, but not the posterior, PG. These findings also corroborate and expand demonstrated associations between the influx of pubertal hormones during adolescence and larger whole PG volumes (Whittle et al., 2020; Wong et al., 2014) by providing further support for the specialized role that the anterior PG plays in pubertal processes and risk for psychopathology. Thus, findings reported here underscore the importance of examining the anterior and posterior PG lobes separately given their functionally distinct roles, particularly when studies are designed to look at puberty-related effects. One possible mechanism explaining the reported patterns could be that increased DHEA during adrenarche promotes enlargement of the PG, a proposed general indicator of HPA-axis activity, which may create amplified risk for mental health symptoms (Farrow et al., 2020; Ganella et al., 2015; Kaess et al., 2018; Whittle et al., 2012). Indeed, there is extensive support for links between patterns of HPA-axis reactivity and risk for psychopathology during adolescence (Colich et al., 2015; Kuhlman et al., 2018), but this work has primarily focused on salivary cortisol with limited links to either structural or functional brain measures. This interpretation needs further investigation with respect to how the timing, tempo, and hormonal changes across the pubertal window explain variability in HPA-axis functioning and anterior PG lobe growth in ways that either propagate or mitigate psychopathology symptoms. To disentangle how different HPA-related hormones influence these processes, more research expressly investigating the interplay between stress and adrenarcheal hormones (e.g., cortisol, DHEA, and their ratios; (Farooqi et al., 2018)) would garner greater understanding into how these complex processes influence puberty-related increases in mental health risk. Ideally, such studies would leverage longitudinal data, tracking dynamic changes in adrenarche indicators (i.e., hormonal and physical) and stress markers (e.g., cortisol levels, inflammatory markers), as well as concomitant changes in anterior PG morphology. Such an approach would contribute greater insight into how patterns of HPA-axis function and pubertal development propagate neuroendocrinological risk for psychopathology.

In line with more transdiagnostic approaches to characterizing developmental psychopathology (Choate et al., 2023), this study is among the first to show anterior PG-mediated pubertal hormone effects in a transdiagnostic symptom profile. To date, extant work has provided much needed insight into associations between overall PG volume and diagnosed psychopathology (e.g., PTSD: Thomas and De Bellis, 2004; depression/bipolar: MacMaster et al., 2008; MacMaster and Kusumakar, 2004; OCD: Atmaca et al., 2009; MacMaster et al., 2006). However, these findings are quite mixed, with some reporting larger or smaller PG volumes, even within the same diagnostic category (e.g., depression; Kessing et al., 2011). Moreover, others have not found associations between diagnostic status and PG volume (Chen et al., 2004; Lorenzetti et al., 2009; Sassi et al., 2001; Schär et al., 2022), which is likely due to a constellation of methodological differences (Anastassiadis et al., 2019), including heterogeneity or sub-phenotypes within one diagnostic category and/or comorbidities. Taken together, the discrepancies in findings may indicate that binary diagnostic categories are not necessarily as informative as more dimensional approaches aimed at capturing neurobiological variability related to common features shared across disorders affecting similar neural circuitry (Cuthbert, 2014; Cuthbert and Insel, 2013). Indeed, there is utility in examining transdiagnostic factors during development, as mental health disorders may not yet be evident or meet diagnostic criteria. Taking this approach has yielded crucial insights into a range of specific transdiagnostic behaviors that portend psychopathology over the short- and long-term (Klein et al., 2022; McLaughlin et al., 2014; Weissman et al., 2020). Dysregulation, in particular, has been shown to be highly predictive of more severe, unremitting psychopathology that encompasses both internalizing and externalizing features (Althoff and Ametti, 2021). In fact, dysregulation symptoms have been proposed as a generalized at-risk phenotype for pronounced risk of psychopathology and persistently poorer outcomes in adulthood, distinct from youth who evince either internalizing or externalizing symptom profiles (Blok et al., 2022; Oerlemans et al., 2020). The current findings add to a somewhat mixed literature, but are largely in line with other findings showing PG-mediated puberty- and DHEA-related increases in internalizing symptomology during development (Murray et al., 2016; Whittle et al., 2012). This study is among the first to examine a transdiagnostic symptom profile that also incorporates externalizing symptoms in this context, but more work is needed to continue to disentangle how particular transdiagnostic symptomologies (e.g., irritability, rumination, emotional awareness) can be explained by different pubertal hormones that convey changes in PG morphology.

Notably, the finding that greater DHEA levels predicted longitudinal decreases in dysregulation symptomology was unexpected, but consistent with existing literature documenting buffering effects of elevated DHEA against mental health symptoms (Mocking et al., 2015). In addition, baseline levels of DHEA did not correspond with baseline levels of dysregulation, suggesting that these associations were best captured with a developmental approach in this sample. It is not intuitive that greater hormonal levels would relate to decreasing symptoms, as there is longstanding evidence that pubertal development is a risk factor for psychopathology. However, work documenting links between puberty and psychopathology has overwhelmingly been focused on gonadarche, and not necessarily adrenarcheal processes per se (Byrne et al., 2017; Pfeifer and Allen, 2021). Therefore, there is still much to uncover with respect to how DHEA levels during the pubertal window, and adrenarcheal processes more broadly, relate to mental health symptoms. Importantly, existing work distinguishes acute and basal DHEA levels, particularly in response to stressors, with documented increases in DHEA in adolescents following an acute stressor (Marceau et al., 2012; Shirtcliff et al., 2007). Conversely, systematically low levels of DHEA in adolescents with internalizing or externalizing symptoms compared to controls have also been documented (Kamin and Kertes, 2017). This suggests that attenuated DHEA levels may correspond with altered HPA-axis physiology in the context of chronic stressors (Jiang et al., 2017), which may be attributable to cortisol and DHEA demonstrating opposing actions in terms of stress-responsivity, with DHEA exhibiting anti-glucocorticoid properties (Pinto et al., 2015). The addition of the PG as a mediator adds to this literature by lending insight to a potential pubertal mechanism that is distinguishable from other potential processes (e.g., stress-related symptomology), although this should be more thoroughly disentangled in future work. That is, research specifically interrogating the involvement of acute and chronic stress as well as pubertal processes in PG growth and the emergence of mental health symptoms is necessary. Exploration of the unique effects of DHEA/DHEA-s, the ratio of DHEA to cortisol, in concert with characterization of physical changes associated with adrenarche would further disentangle the intersection of stress- and puberty-related hormonal cascades that contribute to PG growth and adolescent-specific upticks in psychopathology.

Although the present investigation contributes to a small but growing literature on pubertal hormone effects on PG size and mental health, this study is not without limitations. Specifically, changes in DHEA were not tracked longitudinally in the current study, which could provide crucial information on correspondence between DHEA levels, PG volume, and mental health symptomology. This would allow for more causal interpretations as well as discovery of potential feedback loops between hormone levels and mental health symptoms in youth. In this vein, it will be critical for future work to address how DHEA/DHEA-S levels correspond with physical indicators of adrenarche to understand if more comprehensive measures of adrenarche can explain variability in mental health symptomology and corresponding neural development during adolescence. In addition, participants in this study did not have clinical levels of psychopathology upon enrollment and exhibited subclinical symptom levels. Thus, future work should seek to replicate and extend these effects in higher-risk samples. However, we argue that there is a need to characterize the full spectrum of symptomology, especially in developmental samples, where subclinical or preclinical symptoms could set the stage for diagnostic levels of psychopathology in the long term. Finally, although DHEA levels were assessed at roughly the same time of day in our participants, future work should use this as a covariate. It should be noted, however, that DHEA does not typically have an initial awakening “burst” response as other hormones do (e.g., cortisol), and tends to be more stable day-to-day compared to cortisol in adolescent and adult samples (Hucklebridge et al., 2005; Oskis et al., 2012).

Taken together, the present study provides a novel contribution to a body of literature documenting the role of pubertal hormones and the PG in developmental psychopathology. Using structural equation modeling, we report that with higher DHEA levels, specific mediating effects of the anterior, but not the posterior, PG volume confer risk for increasing dysregulation symptoms across time. This adds important nuance to the existing literature, which has largely focused on estimating whole PG volumes without consideration of the distinct functional roles of the anterior and posterior lobes. Moreover, this work underscores the utility of examining transdiagnostic factors, such as dysregulation, which carry known predictive ability for long-term psychopathology. Identifying how rising levels of DHEA influence an apex neural region within the HPA-axis and changing symptomology is a crucial step in characterizing what role adrenarche plays in adolescent mental health functioning.

## Acknowledgements

This work was supported by the National Science Foundation (#1539067 to TWW, YPW, JMS, and VDC, #2112455 to VDC), the National Institutes of Health (R01-MH121101, R01-MH116782, and R01-MH118013 to TWW; P20-GM144641 to GP and TWW; R01-EB020407 and R01-MH118695 to VDC), and At Ease, USA. Funding agencies had no part in the study design or the writing of this report. All salivary assays were performed at the University of Nebraska-Lincoln Salivary Bioscience Laboratory. We thank Dr. Jessica L. Calvi for her insight and assistance with this aspect of the study. The authors have no conflicts of interest to declare. The data that support the findings of this study are available from the corresponding author upon reasonable request.

