## Supplement for "Anterior pituitary gland volume mediates associations between pubertal hormones and changes in transdiagnostic symptoms in youth"

Table S1. Correlations between all study variables

| | 1. | 2. | 3. | 4. | 5. | 6. | 7. | 8. | 9. |
|---|---|---|---|---|---|---|---|---|---|
| 1. Age | - | | | | | | | | |
| 2. Sex | -.01 | - | | | | | | | |
| 3. Site | .02 | -.13 | - | | | | | | |
| 4. ICV | .11 | **-.53**** | -.02 | - | | | | | |
| 5. DHEA | **.31**** | .01 | .11 | -.08 | - | | | | |
| 6. Dysregulation (baseline) | -.01 | -.02 | .06 | -.14 | .01 | - | | | |
| 7. Δ Dysregulation | .08 | -.002 | .04 | .02 | -.03 | -.20 | - | | |
| 8. Anterior pituitary | **.56**** | **.33**** | **-.20*** | .01 | **.27**** | -.06 | .14 | - | |
| 9. Posterior pituitary | **.24*** | -.04 | **-.55**** | **.21*** | -.14 | -.14 | -.05 | **.19*** | - |

Note. Significant correlations are bolded; ***p* < .01, **p* < .05. ICV = intracranial volume.

Table S2. Standardized and unstandardized path estimates for direct effects

| | | Path Estimates | | | | | |
|---|---|---|---|---|---|---|---|
| IV | DV | β | SE | *p*-value | *b* | SE | *p*-value |
| **Age** | Anterior pituitary | **0.40** | **0.08** | **<.001** | **0.34** | **0.07** | **<.001** |
| **Sex** | | **0.38** | **0.08** | **<.001** | **1.20** | **0.25** | **<.001** |
| **Site** | | **-0.18** | **0.07** | **.01** | **-0.58** | **0.22** | **.01** |
| ICV | | 0.14 | 0.08 | .07 | 0.15 | 0.08 | .08 |
| **DHEA** | | **0.28** | **0.08** | **<.001** | **0.52** | **0.15** | **<.001** |
| Dysregulation (baseline) | | 0.01 | 0.07 | .83 | 0.02 | 0.11 | .83 |
| **Age** | Posterior pituitary | **0.28** | **0.09** | **.002** | **0.05** | **0.02** | **.002** |
| Sex | | -0.05 | 0.09 | .62 | -0.03 | 0.07 | .61 |
| **Site** | | **-0.52** | **0.07** | **<.001** | **-0.38** | **0.06** | **<.001** |
| ICV | | | | | 0.03 | 0.02 | .16 |
| DHEA | | | | | -0.04 | 0.04 | .30 |
| Dysregulation (baseline) | | | | | -0.03 | 0.03 | .31 |
| Age | Δ Dysregulation | 0.08 | 0.15 | .61 | 0.03 | 0.06 | .61 |
| Sex | | -0.03 | 0.12 | .81 | -0.04 | 0.17 | .81 |
| Site | | 0.14 | 0.14 | .33 | 0.20 | 0.20 | .33 |
| **Dysregulation (baseline)** | | **-0.23** | **0.11** | **.04** | **-0.17** | **0.08** | **.04** |
| **DHEA** | | **-0.31** | **0.13** | **.02** | **-0.25** | **0.11** | **.02** |
| **Anterior pituitary** | | **0.35** | **0.16** | **.03** | **0.16** | **0.07** | **.03** |
| Posterior pituitary | | -0.05 | 0.14 | .72 | -0.10 | 0.27 | .72 |
| **Age** | DHEA | **0.53** | **0.07** | **<.001** | **0.24** | **0.04** | **<.001** |
| Sex | | 0.12 | 0.08 | .14 | 0.21 | 0.14 | .14 |
| Site | | 0.14 | 0.08 | .08 | 0.25 | 0.14 | .08 |
| Dysregulation (baseline) | | 0.01 | 0.08 | .91 | 0.01 | 0.07 | .91 |

Note. Bolded results are $p < .05$. These model estimates include the path estimate results from the structural equation model testing mediating effects of anterior and posterior pituitary volume on the association between DHEA levels and changes in dysregulation symptoms. Age, sex, study site, and baseline dysregulation symptoms were included as covariates for all variables in the model and ICV served as a covariate for the pituitary volume variables. CIs = confidence intervals. ICV = intracranial volume.
